# Global Mutational Sweep of SARS-CoV-2: from Chaos to Order

**DOI:** 10.1101/2021.11.16.468834

**Authors:** Xin Wang, Mingda Hu, Yuan Jin, Boqian Wang, Yunxiang Zhao, Long Liang, Junjie Yue, Hongguang Ren

## Abstract

Analysis of large-scale genome sequences demonstrates the mutation of SARS-CoV-2 has been undergoing significant sweeps. Driven by emerging variants, global sweeps are accelerated and purified over time. This may prolong the pandemic with repeating epidemics, presenting challenges to the control and prevention of SARS-CoV-2.

## Main Text

The coronavirus disease 2019 (COVID-19) pandemic, caused by severe acute respiratory syndrome coronavirus 2 (SARS-CoV-2) [1], has been ongoing for more than a year and a half. It has caused over 250 million infections with at least 5 million deaths worldwide as of November 2021. Meanwhile, millions of genome sequences of SARS-CoV-2 have been identified and shared globally. Various mutations have accumulated in the genome of the causative agent since its first identification, resulting in diverse variants of SARS-CoV-2 [2]. As a newly host-jumping virus to human beings, it is almost impossible to predict future new mutations or variants of SARS-CoV-2. However, evolutionary trends can be abstracted from the mutational pattern of large-scale genomes of SARS-CoV-2.

Based on 2,487,499 high-quality SARS-CoV-2 complete genome sequences (see Supplementary_Fasta_ID.csv) from GISAID Website [3], we calculated the nucleotide mutation of each genome in comparison with Wuhan-Hu-1 (GenBank accession number NC_045512). We use the weekly mutation spectrum of the genomes to depict each country and compare their similarities within each two-week window from Feb 24, 2020, to Aug 16, 2021. The similarities are calculated based on both the Cosine similarity (with windows flattened into row vectors) and the Frobenius similarity (the minus Frobenius norm of the matrix difference, see Supplementary Methods). Heat maps of these similarities were generated, along with the stack plots of the proportion of sequences that fall into defined variant groups. Interestingly, although the heat map and the stack plots were generated separately by mutational similarities and infected proportions, a convergent phenomenon was observed between them in the figure (Figure 1).

**Figure 1:**
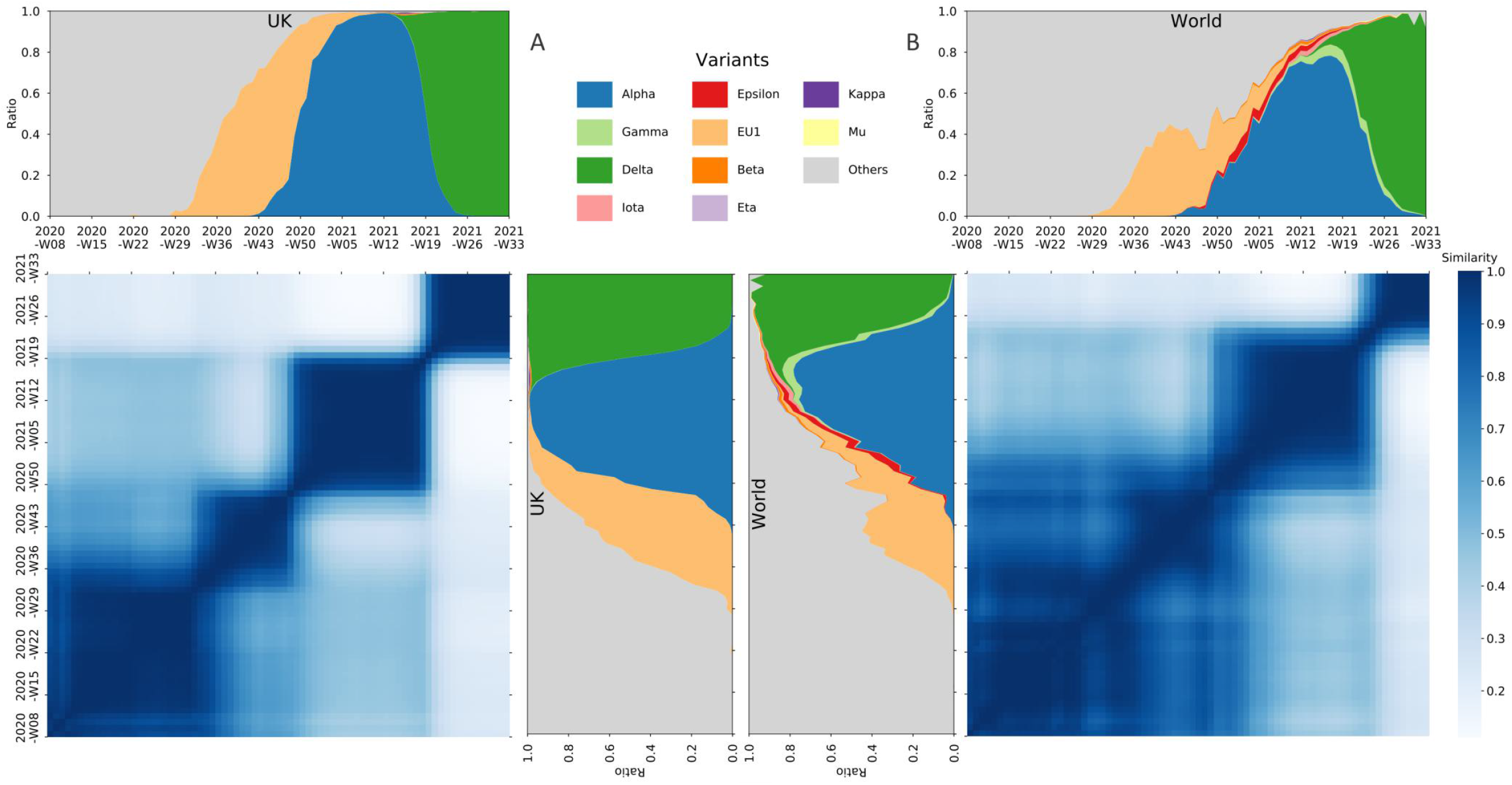
The Cosine similarity of the mutational spectrum of the SARS-CoV-2 genomes within the UK (A) and the whole world (B), along with the stack plots of the proportion of the number of sequences, over time, that fall into defined variant groups.

It has been reported recently that the evolution of SARS-CoV-2 might have been undergoing selective sweeps [4]. While from our observations, the mutation of SARS-CoV-2 genomes has evolved from an early chaotic phase to a state dominated by certain variants. The mutational heat map clearly demonstrates successive dark squares (see Figure 1, Supplementary Figure S1-S10), with each indicating relatively similar and stabilized mutation spectrums during that time period. Each square represents a specific phase during the pandemic, which synchronizes well with the contemporaneous predominant variant in the stack plot. From one phase to the next, the mutation of SARS-CoV-2 undergoes a significant sweeps, in which previous variants (mutation combinations) are swept and replaced by new ones with possible adaptive advantages. Over time, the replacing process for later sweeps may have been accelerated, which can be seen from the sharp borders of later squares in the figure. Furthermore, later squares are darker than earlier ones, suggesting an increasingly genomic homogeneity over phases, which is due to more purified sweeps of variants as the pandemic goes on. The driving forces behind this phenomenon may be related to the enhanced fitness or adaptation of variants to human beings.

We have examined the aforementioned observation in a number of countries with large-scale SARS-CoV-2 sequences separately. Despite the regional differences, the conclusion holds for almost all situations (see Supplementary Figure S1-S10), which all show phases (squares) divided by mutational sweeps. Benefit from abundant genome sequences, the figure of the United Kingdom is considerably representative (see Figure 1A), which shows several clear phases divided by sweeps dominated by typical SARS-CoV-2 variants.

We further compared the mutation spectrums among different regions. Due to the difference in both the control measure and the prevalence of variants, the heat map shows a variety of shapes. Taking comparing the UK and the US as an example (see Supplementary Figure S11-S12), the persistence of *Alpha* variant and *Delta* variant in the UK are longer than those in the US, so the similar mutation spectrum shows rectangles rather than squares. Note the mutation spectrums of these two countries showed less similarities in 2020, indicating the early regional genomic difference between the two countries. Nevertheless, the emergence of *Alpha* and then *Delta* variants in 2021 quickly converged such regional diversity. This implies that the evolution of SARS-CoV-2 has been undergoing selective sweeps both regionally and globally, in which previous local predominant strains can be quickly replaced by imported variants, e.g., *Alpha* and *Delta* variants, which has evolutionary advantages either in transmission or host adaptation, or both.

At the time this manuscript being submitted, the *Delta* variant has almost completed its sweep process throughout the world and become a global dominant variant. New SARS-CoV-2 variants with enhanced fitness will surely emerge in the future to replace the former predominant variant, but the possibility of co-circulating of multiple competing variants is low. It seems that the SARS-CoV-2, after the host-jumping event, has finished the early stages in adaptation to human beings through chaotic mutations and evolved into relatively persistent stabilized adaptations. More or less like the seasonal influenza virus [5], the alternation of epidemic strains of SARS-CoV-2 may become periodic.

Altogether, our study demonstrates that the SARS-CoV-2 has evolved from early chaotic mutations into relatively persistent stabilized adaptations. This ongoing adaptation presents successive phases of the pandemic, along with stronger sweeps and increasingly global homogeneity driven by the continuous emergence of SARS-CoV-2 variants. The completion of stage transition of COVID-19 might substantially prolong the pandemic with repeating epidemics, making it vital to strengthen the surveillance of SARS-CoV-2 globally. Meanwhile, vaccine development and vaccination strategies may should be updated accordingly.

## Supporting information

Supplementary_Fasta_ID

## FUNDING

This work was supported by The National Natural Science Foundation of China [grant number 32070025, 31800136, 82041019].

## Conflict of interest statement

None declared.

## Appendix

### Materials and Methods

We collected 2,487,499 high-quality SARS-CoV-2 complete genome sequences from GISAID Website (c.f. Fasta ID.csv for detailed information). For each genome, the nucleotide mutation is calculated in comparison with Wuhan-Hu-1 (GenBank accession number NC_045512). We study the mutation spectrum of genomes in a given region, focusing on the whole world and four object countries, Brazil, India, the United Kingdom, and the United States. Nucleotide mutations with global occurrence of less than 10,000 are considered infrequent and then abandoned. Additionally, mutations absent in any of the four countries are neither included. Thus, 475 major nucleotide mutations remain for further studies.

We depict each region by its weekly mutation spectrum of genomes, consisting of the weekly proportion of 475 major mutations from Feb 24, 2020, to Aug 16, 2021, namely a period of 78 weeks. A two-week window is screened in the spectrum and the similarity is calculated, using both the Cosine similarity and the Frobenius similarity.

For the Cosine similarity, the matrix of each window is flattened into a row vector, then the Cosine similarity between flattened vectors ***a*** and ***b*** is calculated as follows.

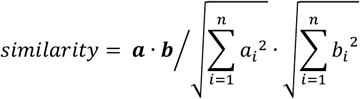

For matrices of two windows, ***A, B*** ∈ ℝ^*m*×*n*^, we define the Frobenius similarity between windows by the minus Frobenius norm of the difference between matrices ***A*** and ***B***.

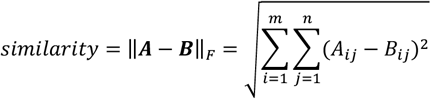

Heat maps are generated from these similarities between spectrums. For each region, we also analyze the SARS-CoV-2 genome sequences and draw the stack plots of the proportion of defined variant groups. The correspondence between heat maps and contemporaneous variant groups in stack plots is conducive to our studies.

**Supplementary Figure S1.**
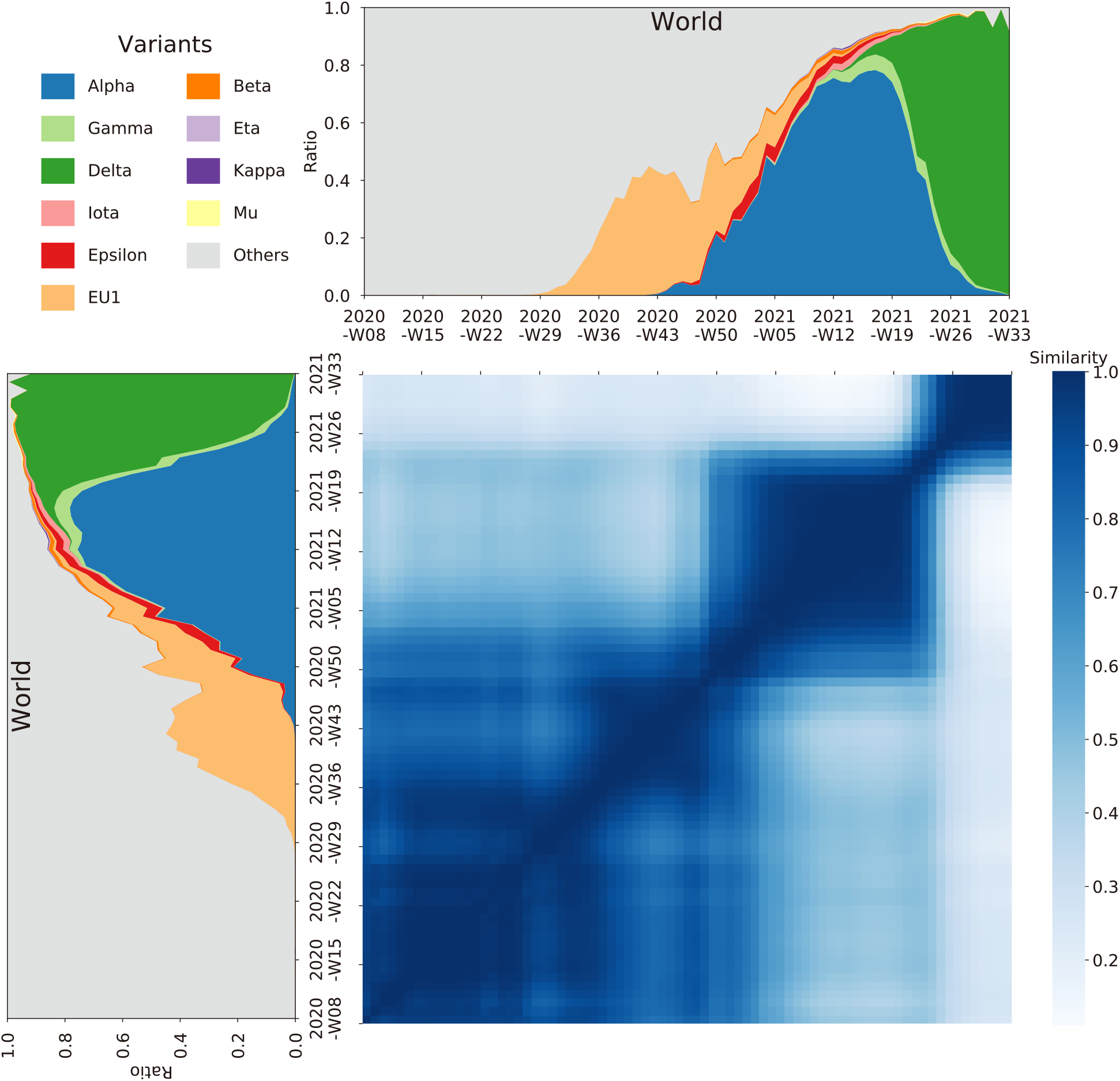
The Cosine similarity of the mutational spectrum of the SARS-CoV-2 genomes within the whole world.

**Supplementary Figure S2.**
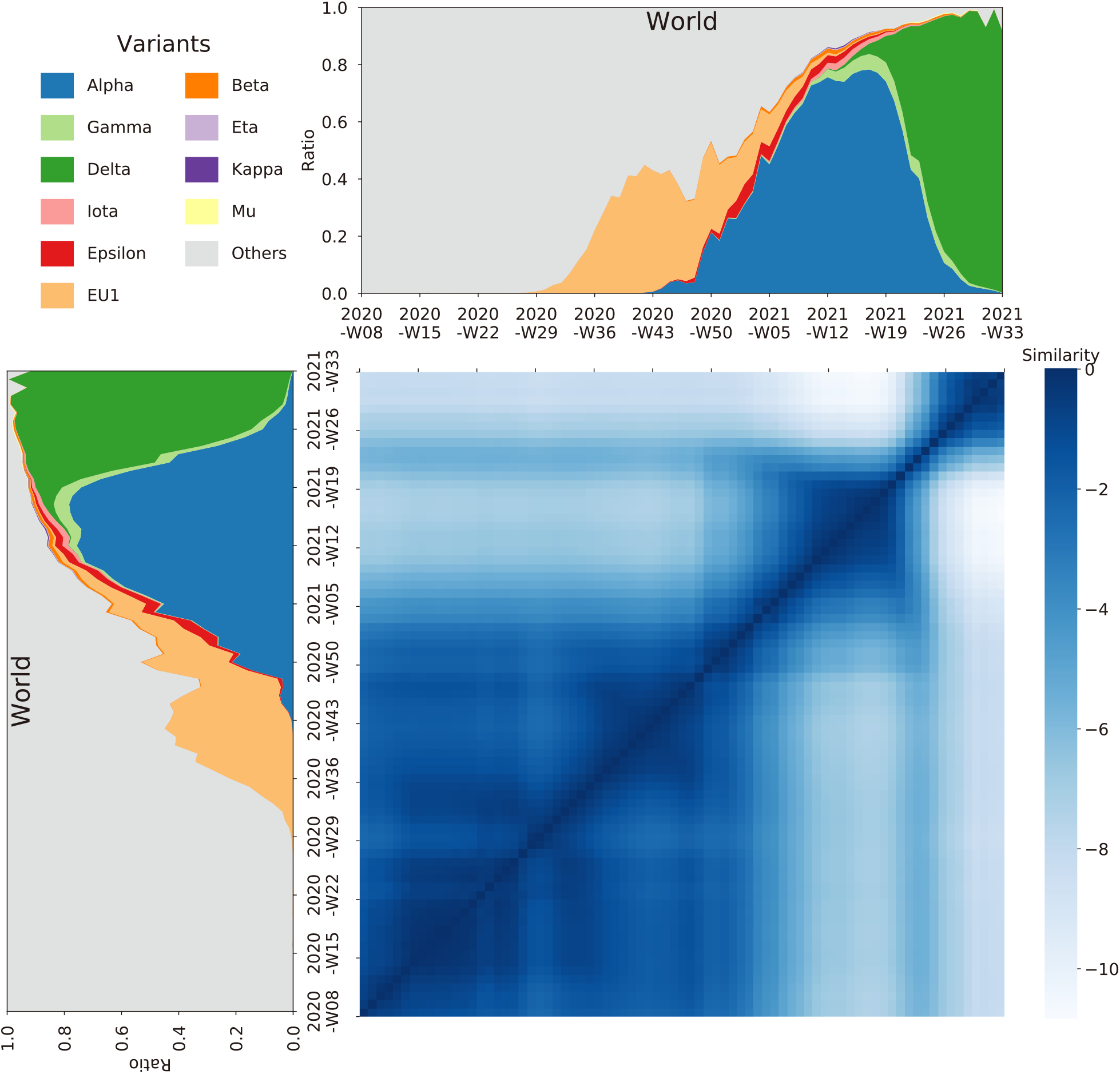
The Frobenius similarity of the mutational spectrum of the SARS-CoV-2 genomes within the whole world.

**Supplementary Figure S3.**
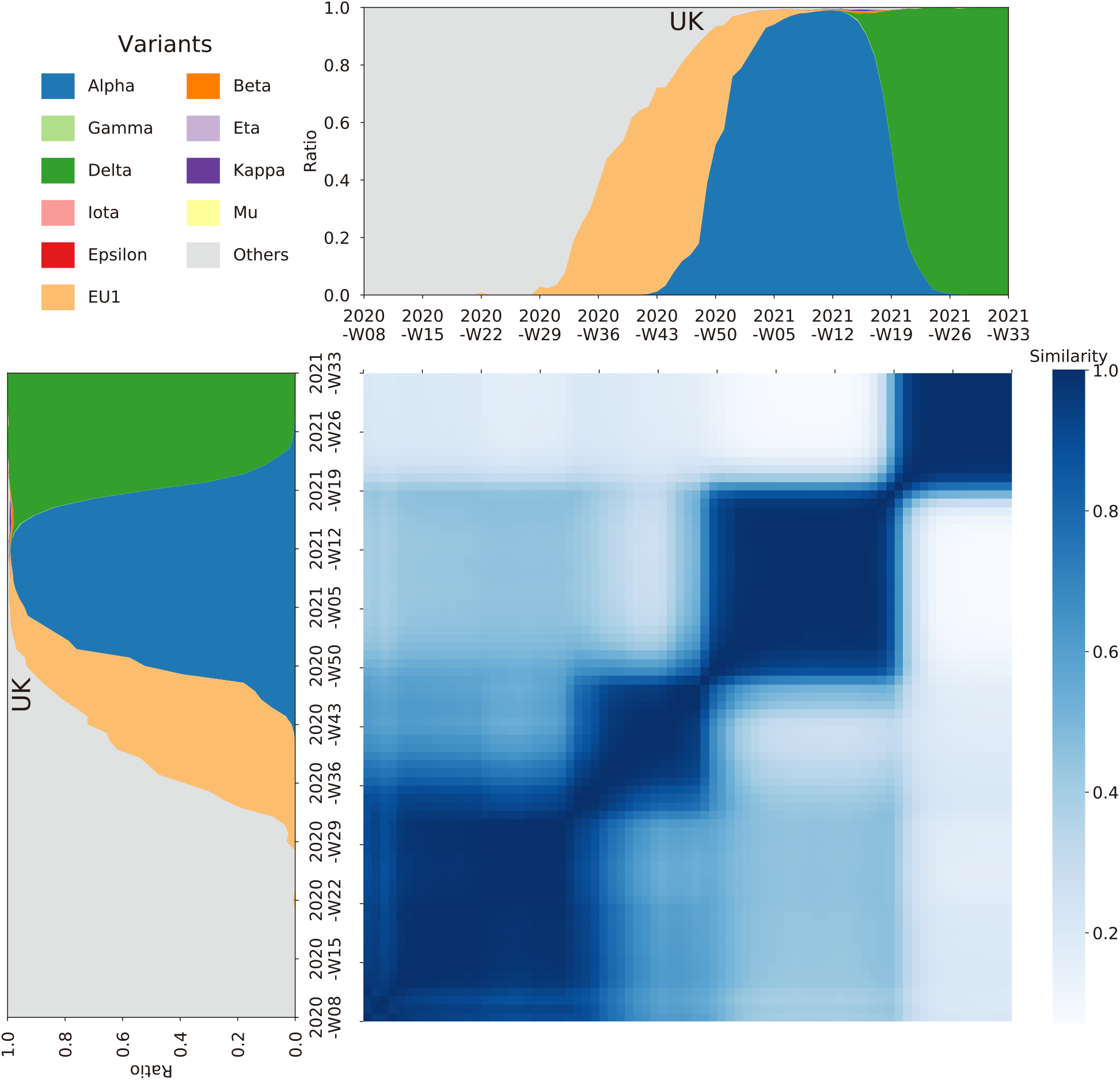
The Cosine similarity of the mutational spectrum of the SARS-CoV-2 genomes within the UK.

**Supplementary Figure S4.**
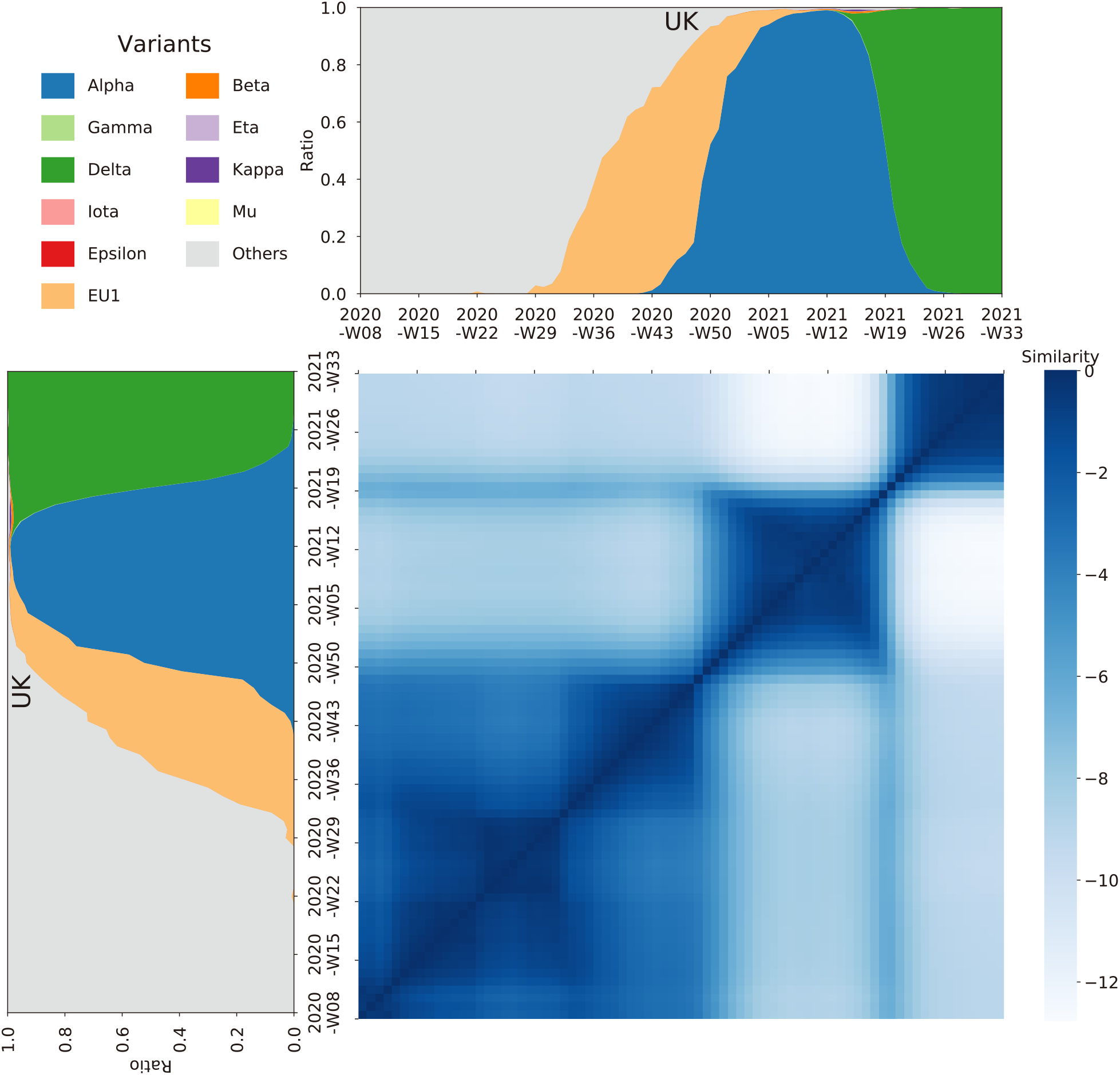
The Frobenius similarity of the mutational spectrum of the SARS-CoV-2 genomes within the UK.

**Supplementary Figure S5.**
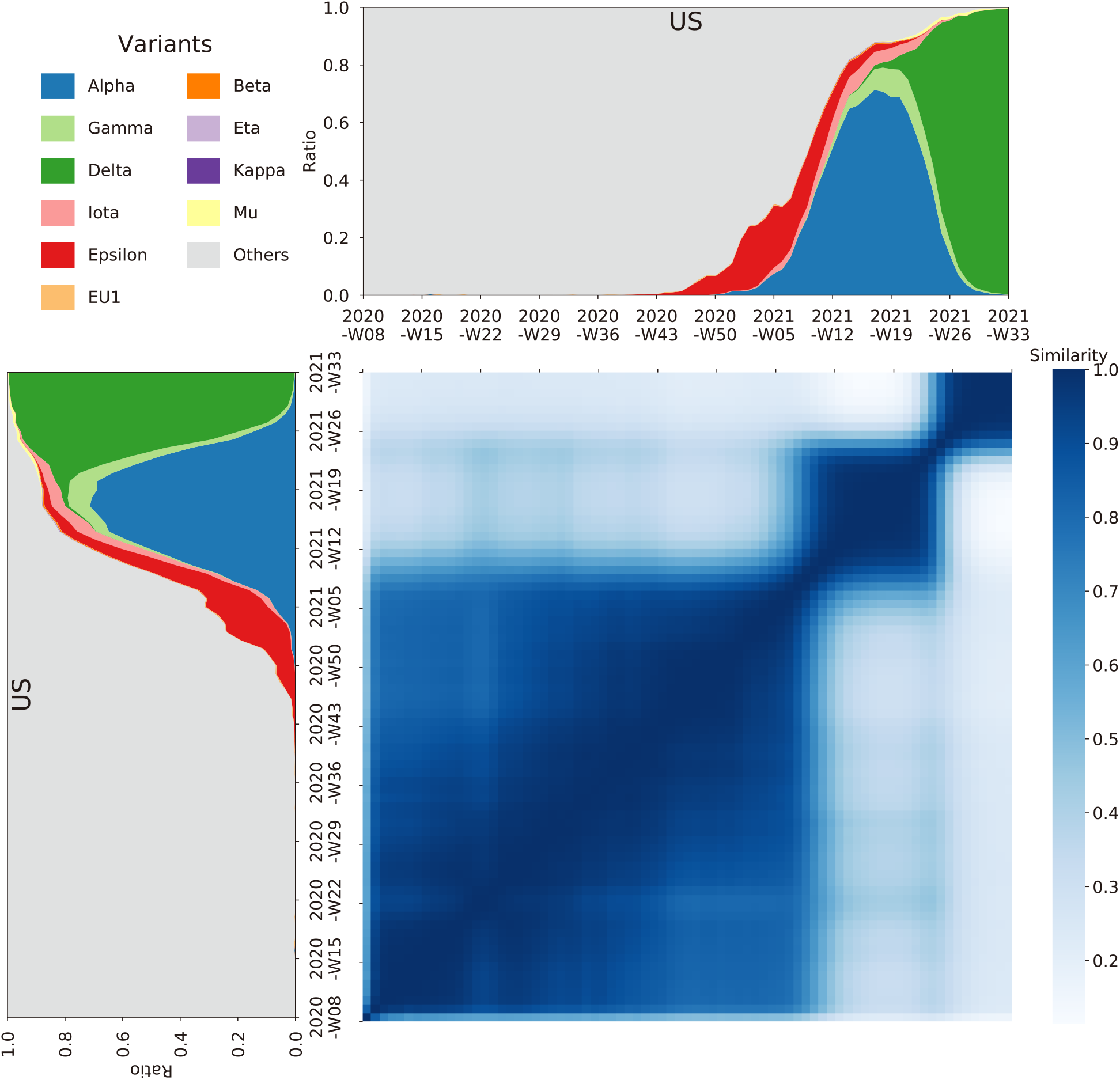
The Cosine similarity of the mutational spectrum of the SARS-CoV-2 genomes within the US.

**Supplementary Figure S6.**
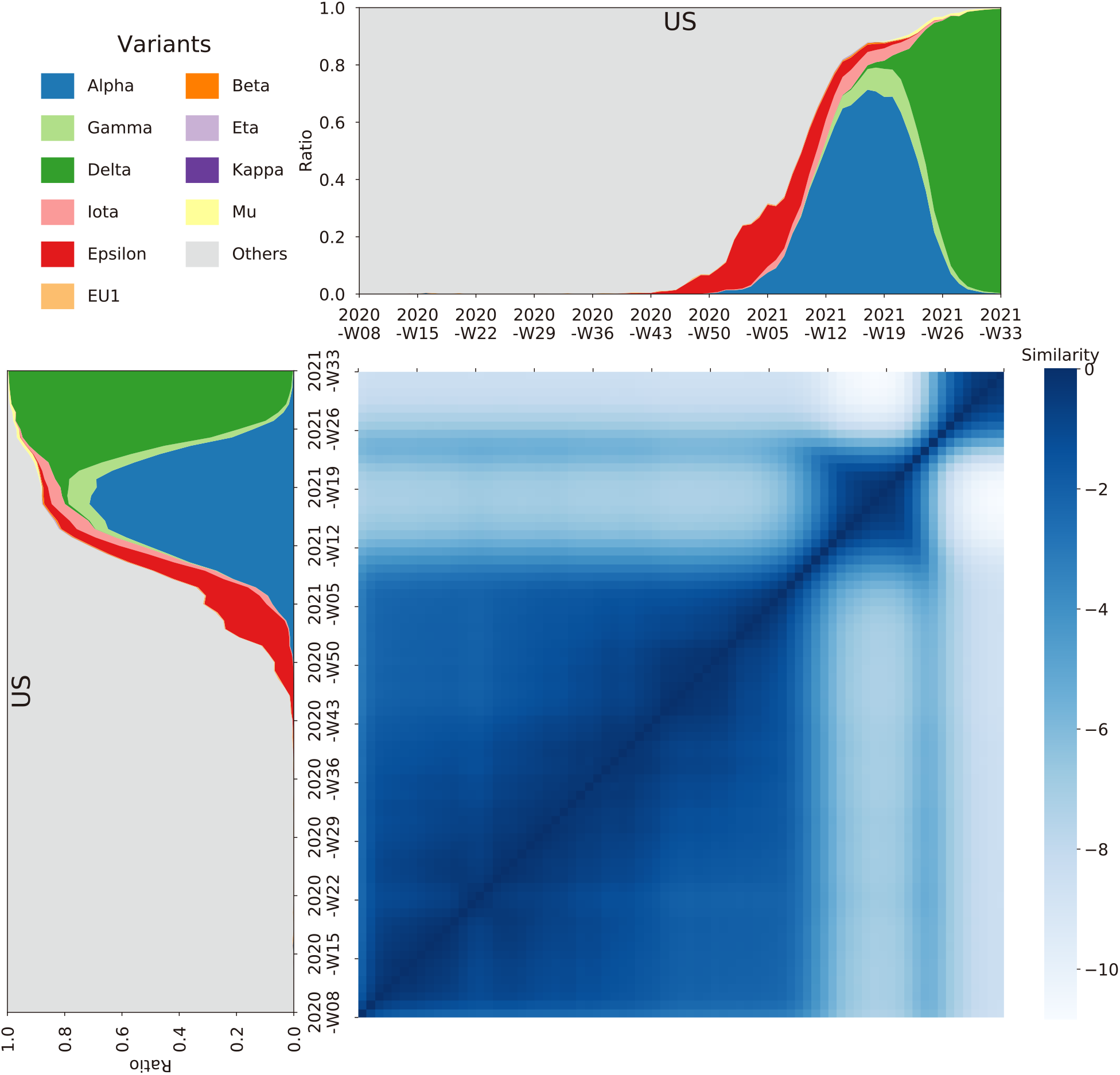
The Frobenius similarity of the mutational spectrum of the SARS-CoV-2 genomes within the US.

**Supplementary Figure S7.**
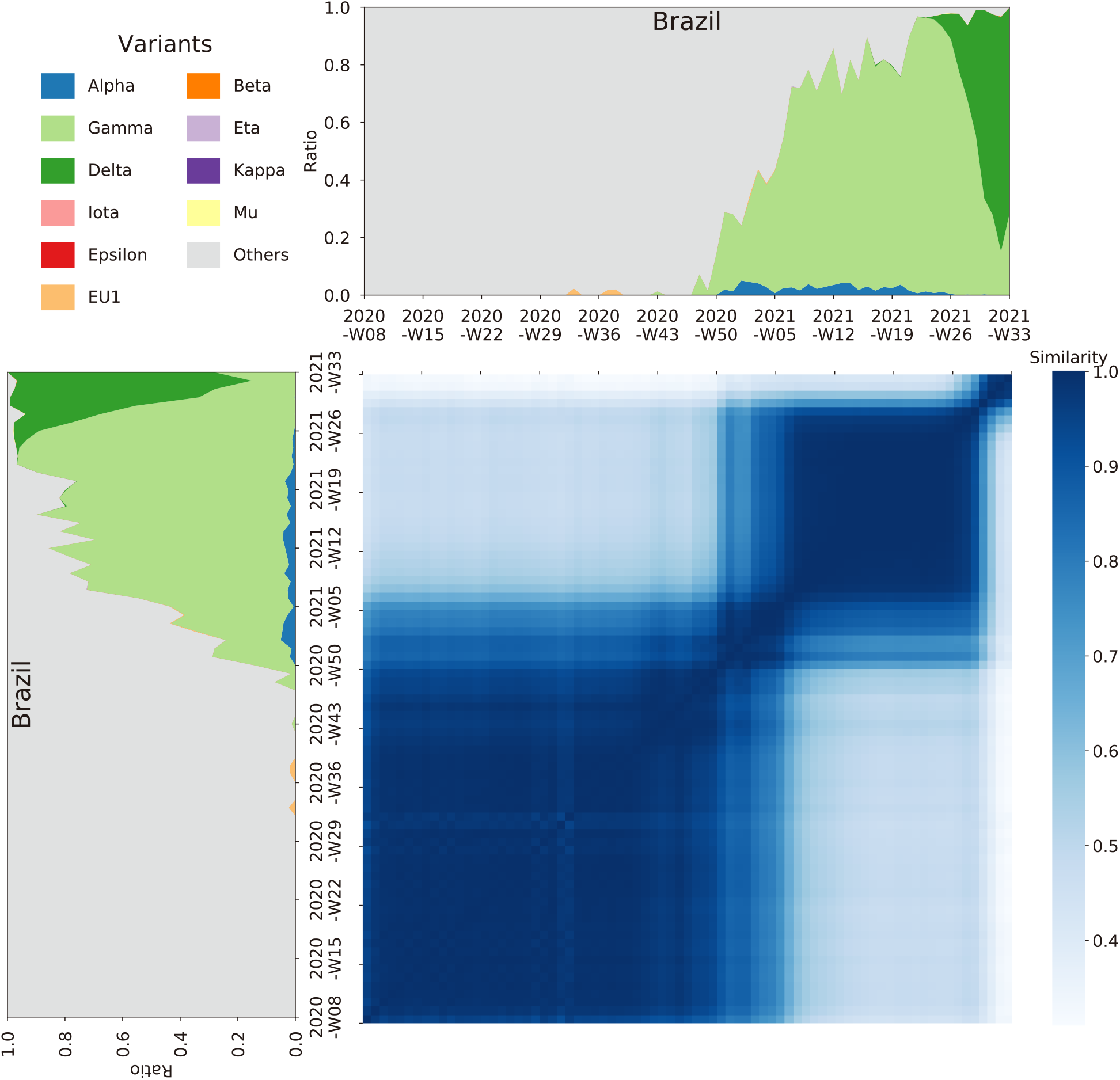
The Cosine similarity of the mutational spectrum of the SARS-CoV-2 genomes within Brazil.

**Supplementary Figure S8.**
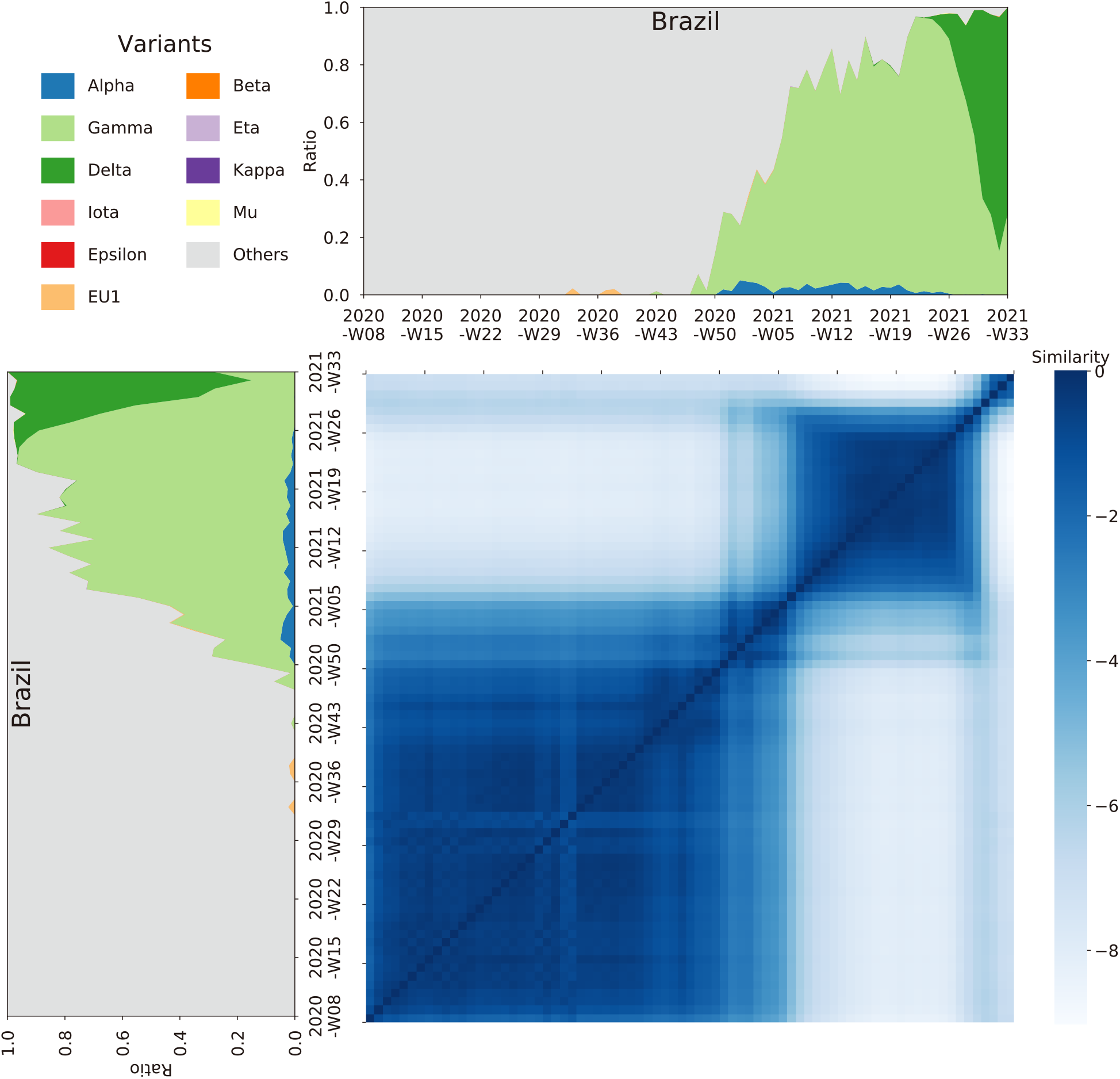
The Frobenius similarity of the mutational spectrum of the SARS-CoV-2 genomes within Brazil.

**Supplementary Figure S9.**
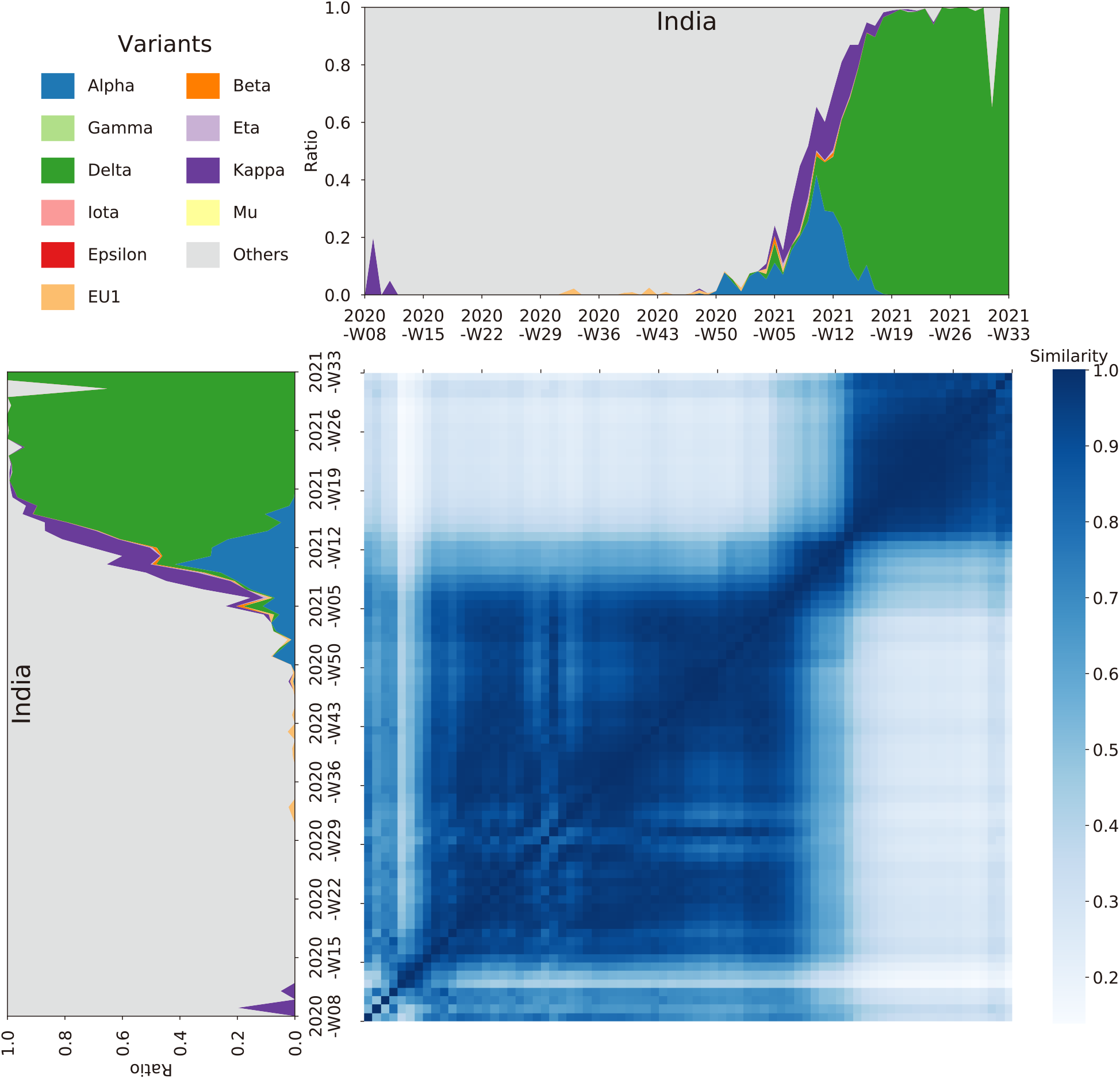
The Cosine similarity of the mutational spectrum of the SARS-CoV-2 genomes within India.

**Supplementary Figure S10.**
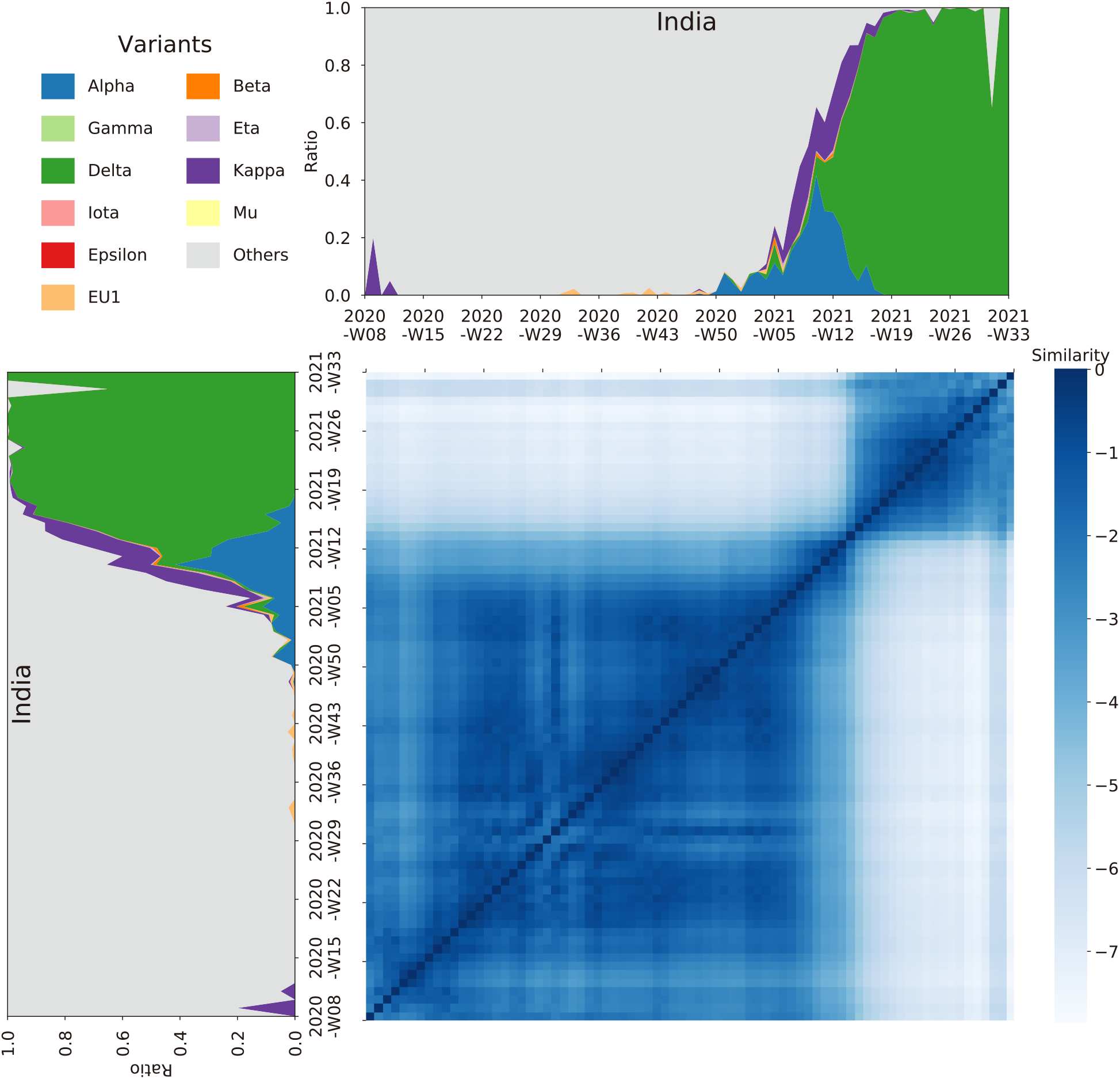
The Frobenius similarity of the mutational spectrum of the SARS-CoV-2 genomes within India.

**Supplementary Figure S11.**
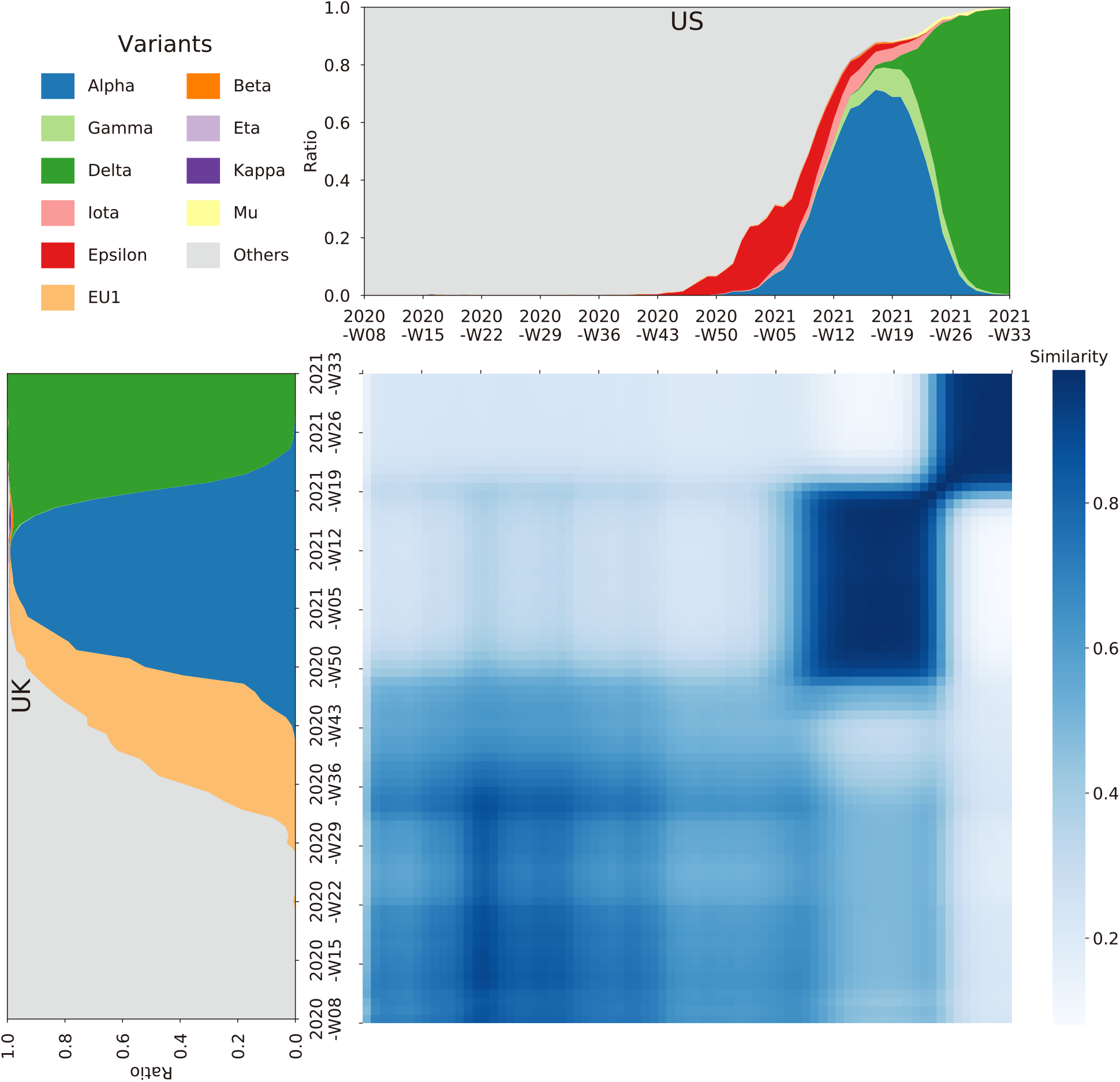
The Cosine similarity of the mutational spectrum of the SARS-CoV-2 genomes between the UK and the US.

**Supplementary Figure S12.**
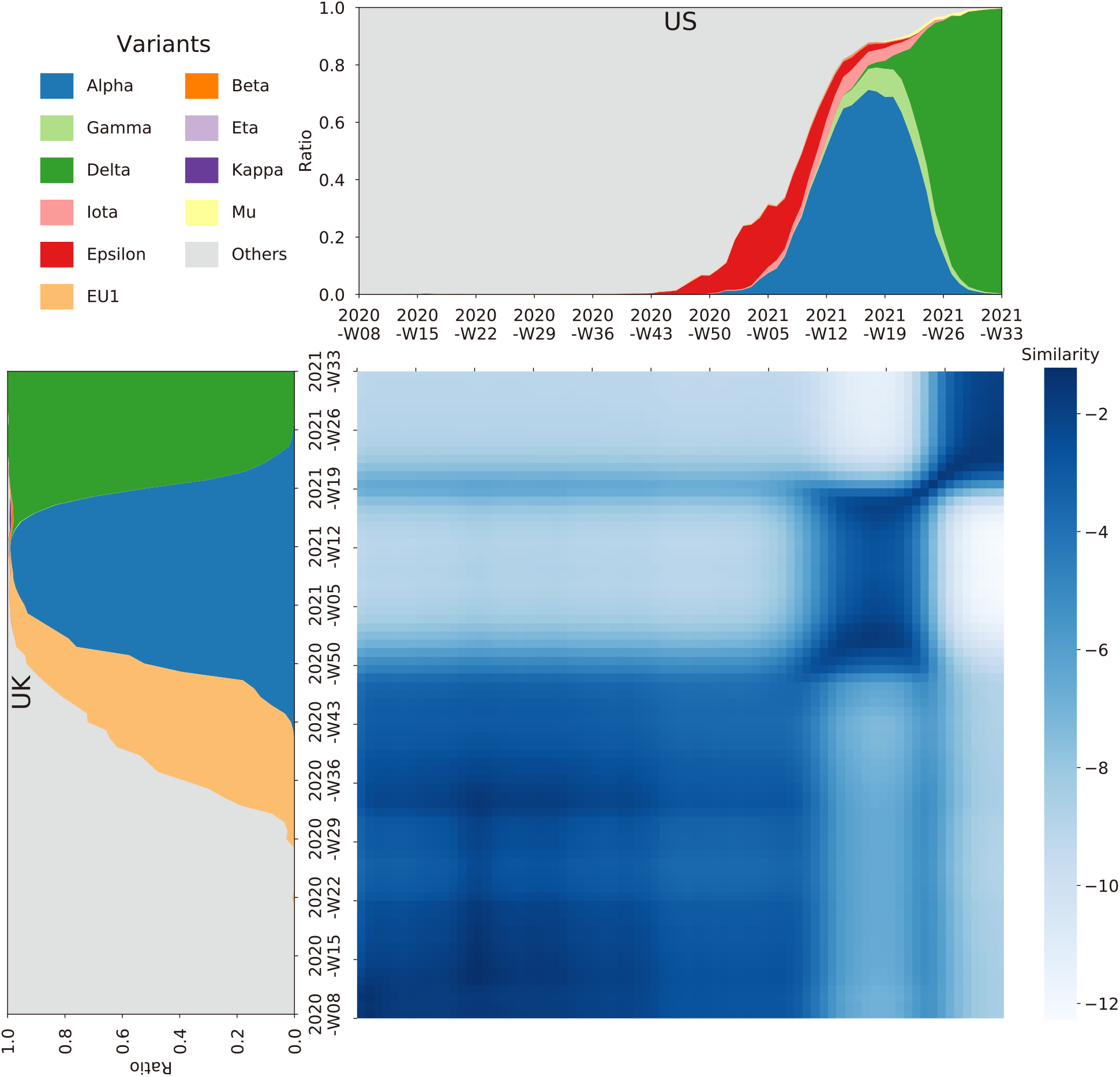
The Frobenius similarity of the mutational spectrum of the SARS-CoV-2 genomes between the UK and the US.

### Genome Sequence Availability

We downloaded SARS-CoV-2 genome sequences as of Sep 08, 2020, from GISAID Website1. Only high-quality complete sequences are retained and thus we obtained 2,487,499 genome sequences of SARS-CoV-2. Please c.f. Supplementary_Fasta_ID.csv for detailed information of those genome sequences.

## Notes

### Competing Interest Statement

The authors have declared no competing interest.

## REFERENCES

1. Zhou P, Yang XL and Wang XG et al. A pneumonia outbreak associated with a new coronavirus of probable bat origin. Nature. 2020;579:270–273.

2. Harvey WT, Carabelli AM, Jackson B et al. SARS-CoV-2 variants, spike mutations and immune escape. Nature Review Microbiology. 2021;19:409–424.

3. Shu Y and McCauley J. GISAID: Global initiative on sharing all influenza data - from vision to reality. Euro Surveill. 2017;22(13).

4. Kang L, He G and Sharp AK et al. A selective sweep in the Spike gene has driven SARS-CoV-2 human adaptation. Cell. 2021;184(17): 4392-4400.e4394.

5. Petersen E, Koopmans M and Go U et al. Comparing SARS-CoV-2 with SARS-CoV and influenza pandemics. The Lancet Infectious Diseases. 2020;20(9):e238–e44.

## References

1. Shu Y and McCauley J. GISAID: Global initiative on sharing all influenza data - from vision to reality. Euro Surveill. 2017;22(13).

